# Two SecA/SecY Systems with Distinct Roles in the Ecological Adaptations of Bacteria

**DOI:** 10.1101/202358

**Authors:** Xiaowei Jiang, Mario A. Fares

## Abstract

Bacteria interact with their environment through the secretion of a specific set of proteins (known as secretome) through various secretion systems. Molecular modifications of these secretion systems may lead to the emergence of new bacterial-environment interactions, although this remains unexplored. In this study we investigate the possible link between molecular and functional changes in secretion proteins and the ecological diversity of bacteria. We studied functional modifications in secretion proteins by identifying events of functional evolutionary divergence—that is, changes at the molecular level that have driven changes of protein’s function. We present data supporting that these functional diversifications occurred in essential secretion proteins in bacteria. In particular, functional divergence of the two most important secretion proteins SecA and SecY in pathogenic bacteria suggests that molecular changes at these proteins are responsible for their adaptations to the host. Functional divergence has mainly occurred at protein domains involved in ATP hydrolysis in SecA and membrane pore formation in SecY. This divergence is stronger in pathogenic bacteria for protein copies resulting from the duplication of SecA/SecY, known as SecA2/SecY2. In concert with these results, we find that the secretome of bacteria with the strongest functional divergence is enriched for proteins specialized in the interaction with specific environments. We unravel evolutionary signatures that link mutations at secretion proteins to the ecological diversification of bacteria.

## Introduction

Bacteria interact with specific environments by translocating a specialised set of proteins across the cell membrane. Approximately one quarter to a third of bacterial proteins are initially synthesized in the cytosol and subsequently targeted to either the cell membrane or to the extra-cellular space of the cell [1]. To translocate proteins across the membrane, bacteria use two major secretion systems, the general secretion (Sec) pathway and the Sec-independent Twin-arginine translocation (Tat) pathway. The Sec pathway translocates proteins in an unfolded state (pre-proteins), while the Tat pathway translocates fully or partially folded proteins. Correct protein translocation relies on N-terminal end signal peptide of the substrate. In bacteria, the Sec system has two independent pathways: the post-translational and the co-translational pathways. In both pathways, pre-proteins are first led to the cytoplasmic membrane across which they are translocated by an evolutionarily conserved heterotrimeric protein complex channel (SecYEG).

In the post-translational pathway, synthesized pre-proteins in the cytosol bind to SecB and/or SecA to be transported to the plasma membrane. Binding of SecB has been shown to be essential to prevent the pre-protein premature folding prior to membrane translocation [2]. In gram-negative bacteria, SecB-pre-protein complex interacts and transfers the pre-protein to SecA. *Escherichia coli* lacks SecB, while SecA seems to complement the activity of SecB [3,4].

In the co-translational pathway, SRP (signal recognition particle) binds to the polypeptide emerging from the ribosome and directs the ribosome-nascent chain complex to the plasma membrane, where the complex binds to the SRP receptor FtsY. Subsequent to this, the emerging polypeptide passes through the SecY translocation channel [5]. In archaea, there is evidence that some proteins can be translocated post-translationally, but archaea lack SecB and SecA, hence post-translational translocation remains to be elucidated [6].

SecA and SecY, that are part of the complex SecYEG-SecA, have been extensively studied as they are directly responsible for the translocation of a large fraction of pre-proteins through the membrane [7,8]. The 100 Kda protein SecA, considered to be a motor protein, participates in pre-proteins translocation mediated by its ATPase activity [9]. This protein is conserved among bacteria and some plants [10], and is essential to cell viability [7]. SecA consists of six protein domains (Figure 1B): nucleotide-binding domain 1 (NBD1) and 2 (NBD2), the polypeptide-cross-linking domain (PPXD), the helical scaffold domain (HSD), the helical wing domain (HWD) and c-terminal domain (CTL, not shown in Figure 1B). SecA function occurs through strong conformational changes mediated by ATP binding and hydrolysis at the NBD domain [11]. PPXD, NBD2 and part of HSD domains have been suggested to form a “clamp” for pre-protein peptide binding and translocation [12]. SecY presents a clam shell-like symmetrical arrangement and consists of ten transmembrane (TM) segments (TM1-TM10), which form a gate at the front side and are clamped together at the back by SecE [12] (Figure 2B). The role of SecG, another component of the complex SecYEG-SecA, has been deemed not essential [13], although it may be involved in mediating the interaction of SecA and SecY [14–17].

**Figure 1.**
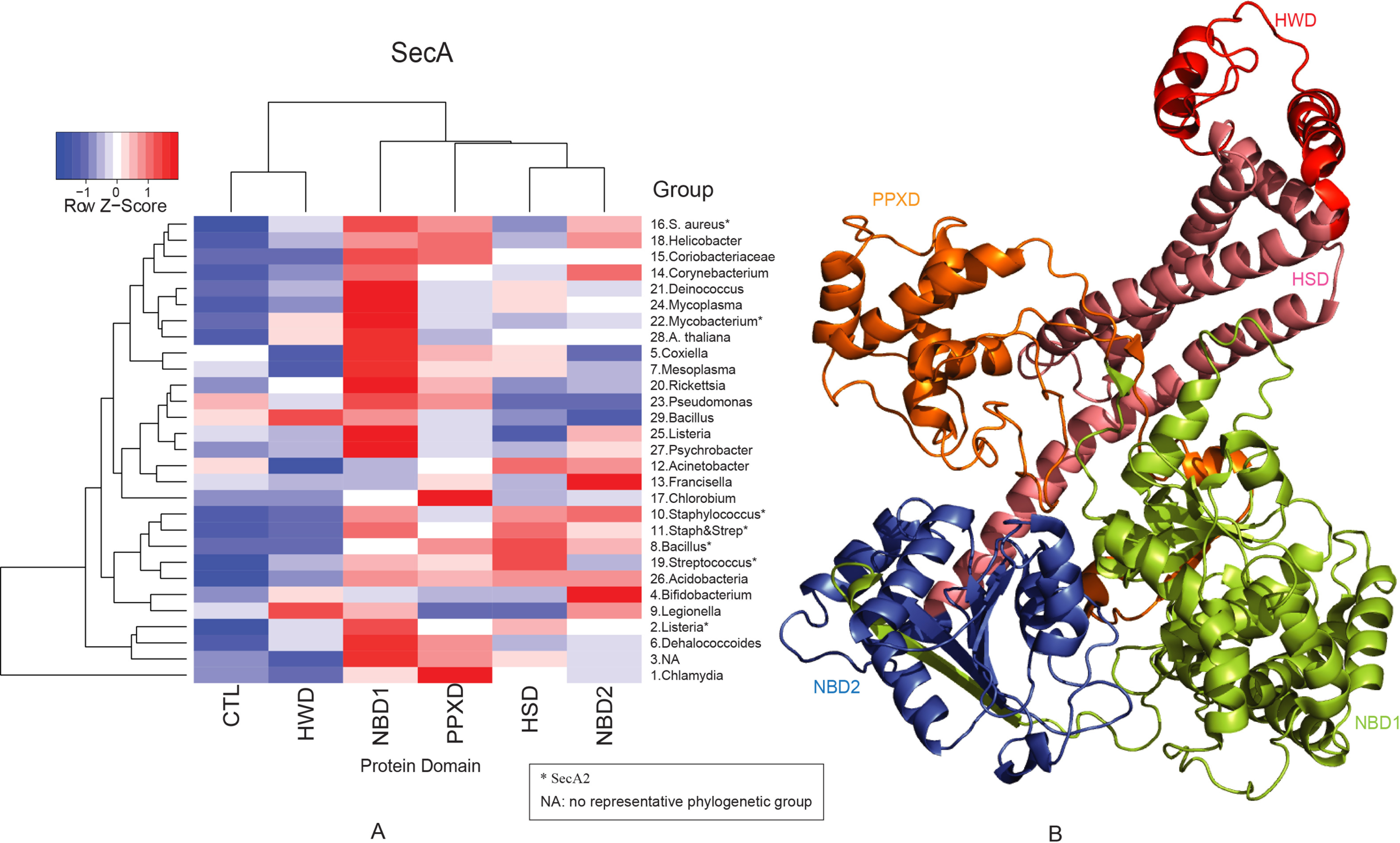
Two-way clustering of the top 10% clades under functional divergence in SecA. (A) Protein domains are clustered according to the number of sites under functional divergence after Z-score standardization, group number represents the rank of number of sites under functional divergence in each analyzed branch, and groups are also clustered. (B) Protein domains are shown in SecA. * denotes functional divergence results for SecA2, representative phylogenetic group in each group is also shown.

Two SecY/SecA systems are found in bacteria, one is termed SecA1/SecY1 (canonical SecA/SecY, interchangeably used in this paper), and the other is called (accessory) SecA2/SecY2, [18]. SecA1 and/or SecY1 perform “housekeeping” functions in bacterial physiology, whereas SecA2 and/or SecY2 are involved in determining species virulence traits [18]. However, the precise role of accessory SecA2/SecY2 system on the overall bacterial physiology, transcription and secretion remains elusive.

SecA2/SecY2 system is frequently present in highly pathogenic and antibiotic resistant bacteria such as *Staphylococcus aureus* (meticillin-resistant, vancomycin-susceptible, vancomycin-intermediate resistance strains) and *Streptococcus pneumoniae*. Although not essential for pathogens survival [18,19], this system has been found to be solely responsible for secreting a set of virulence factors [18] and post-translationally modified glycoproteins [20–29]. Heavily glycosylated pre-proteins are exceptionally long and their translocation may have required major changes in the secretion system, very likely through duplication and functional specialization of SecA2 and SecY2 [30–32]. Although SecA2/SecY2 specialization in translocating other sets of proteins has been previously suggested [33–35], molecular changes responsible for this and their functional consequences remain largely unexplored.

Due to the important role of Sec translocase in bacterial adaptations, mapping molecular and functional evolutionary patterns to ecological specializations is a fundamental aim in evolutionary and microbial biology. To identify these evolutionary patterns, we conducted an analysis of functional evolutionary divergence in key Sec proteins across the bacterial phylogeny. Our results suggest that strong functional divergence in bacterial Sec secretion systems could indeed be associated with their ecological adaptations.

## Materials and Methods

The main objective of this study is to identify evolutionary events in Sec proteins that were responsible for their functional diversification between groups of bacteria and between the canonical SecA/SecY and derived (accessory) SecA2/SecY2 systems. To do so, we have conducted extensive computational analyses to identify patterns of functional divergence (FD) in Sec proteins using a large phylogeny of bacterial species that included microbes with different lifestyles: pathogens, extremophiles and free-living bacteria.

### Sequence selection and alignments

We built multiple sequence alignments of bacterial Sec proteins by retrieving homologous protein sequences for SecA, SecB, SecE, SecG and SecY from KEGG (Kyoto Encyclopedia of Genes and Genomes) pathway database (updated December 1, 2009) using KEGG API (Application Programming Interface) function with Perl programming [36]. The set of homologous protein sequences comprised 944 bacteria, 68 archaea and 10 plants (Table S1, S2, S3 and S4). Pathogenic bacterial strains and their hosts are also identified from KEGG (http://www.genome.jp/files/org2key.xl).

To identify protein sequences of SecA1/SecY1 and SecA2/SecY2, we followed a strategy used before [18]. Briefly, in bacteria with two SecA/SecY homologs, SecA/SecY with a slightly higher sequence similarity to the canonical SecA/SecY of *E. coli* and *Bacillus subtilis* were considered to be SecA1/SecY1, while the SecA/SecY with a lower sequence similarity to SecA and SecY were termed SecA2/SecY2.

To understand how evolutionary changes in SecA were followed by those in SecY and viceversa, we conducted coevolution analyses between SecA and SecY, and SecA2 and SecY2. Multiple sequence alignments were also built to conduct analyses of intermolecular covariation (coevolution, see methods below) between canonical and accessory SecA and SecY. These alignments included 91 bacteria with SecA and SecY, that also presented SecB, and 15 bacterial strains that were found to have both SecA2 and SecY2. To analyze intermolecular coevolution between SecA1 and SecB, we retrieved 83 sequences of each protein, respectively. All protein sequences were aligned by Muscle 3.7 [37] and alignments were manually checked using Jalview [38].

### Phylogenic analysis

RaxML Pthreads version 7.2.4 [39] was used to build the Maximum likelihood (ML) phylogenetic trees after estimation of the appropriate amino acid substitution model by ProtTest version 2.4 [40]. Then, the best ML trees for SecA, SecE, SecG and SecY were computed based on the LG substitution model [41] with GAMMA model of rate heterogeneity and empirical amino acid frequencies.

### Analysis of Functional divergence

Functional divergence is a term used to refer to the divergence, in a group of organisms, of a protein function as a result of the amino acid mutations at that protein. Analysis of functional divergence rests on the assumption that evolutionary conserved amino acid sites are important for protein’s function because their change is deleterious. Functional divergence can be classified in two groups [42] i) FD type I refers to the acquisition of a functional role of an amino acid site in a group of bacteria—that is, the amino acid site becomes conserved within this group resulting from the new selection constraints on the novel function, otherwise this site evolves neutrally; and ii) FD type II according to which the amino acid site is highly conserved in the two different phylogenetically related groups of bacteria but have different functional roles.

We previously devised a method that allows exploring an entire phylogeny for evidence of functional divergence [43,44]. Here, we applied this method to identify groups of bacteria with strong functional divergence. Briefly, the method uses a protein sequence alignment and a phylogenetic tree that includes paralogs and orthologs. It then compares a pair of clades in the tree sharing a common ancestral origin to their closest phylogenetic outgroup. The comparison is performed for each amino acid site in the protein and the strength or likelihood of the amino acid transition state (amino acid mutation) is evaluated using the appropriate BLOSUM matrix (BLOcks of Amino Acid SUbstitution Matrix) [45]. BLOSUM are score matrices that quantify the likelihood of the transition between the 20 amino acids, with positive, 0 and negative scores meaning the transitions are more frequent, as expected and less frequent than expected, respectively. In the comparison between two clades to an outgroup (clade 1 and clade 2), functional divergence in clade 1 would be detected if the BLOSUM scores were positive within clade 1 (more frequent than expected), negative between clade 1 and the outgroup (extreme transitions) and positive between clade 2 and the same outgroup. The functional divergence score is calculated as follow and compared with a normal distribution:

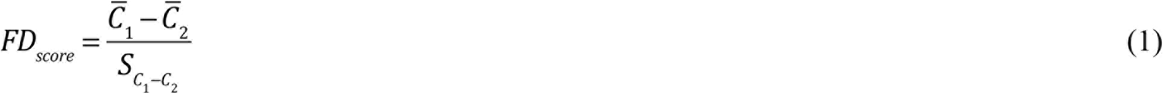

where *C̅*_1,2_ are the mean substitution scores for the transition from clades on either side of the bifurcation in the phylogenetic tree relative to the outgroup and *S*_*C*1-*C*2_ is the standard error for unequal sample sizes with unequal variances.

To identify the patterns of sites under FD among protein domains in SecA/SecYEG proteins, we first ranked each of the clades according to the number of sites under FD in comparison with the rest of clades. We then counted the number of sites under FD within each protein domain for that particular protein and scaled this number according to domain length. Third, we used these numbers to build a data matrix for each protein, where each row represented the FD enrichment profile for the 10% clades most enriched for FD, while columns referred to the FD profiles within each of the protein domains. Finally, we used this data matrix as input for the clustering of the rows and columns in order to search for similarities in the FD profiles of protein domains and/or clades, which was performed using heatmap.2 function in R (http://www.r-project.org/). The heatmap.2 function scales each element in a row in the data matrix based on a normalised Z-score. The score is calculated as follows:

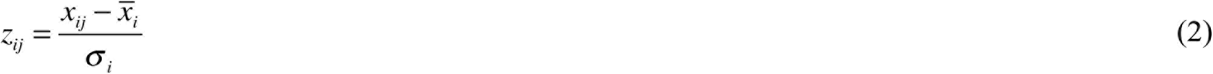

Here, *Z*_*ij*_ is the Z-score calculated for element *x*_*ij*_ in the matrix, *x̅*_*i*_ is the mean of the row *i*, ***σ***_*i*_ is the standard deviation of all the elements in the row *i*.

### Prediction of potential Sec and Tat substrate proteins

To identify potential SecA1/SecY1 dependent substrate proteins, we followed three steps. First, we used SignalP [46] to screen a proteome for potential Sec-dependent secreted proteins (secretome). Second, we used TatP [47] to screen that proteome for potential Tat-dependent secreted proteins. Finally, we remove those proteins predicted by TatP in the SignalP results to minimize the number of false positives. To identify potential SecA2/SecY2 dependent substrate proteins, the reciprocal best-best FASTA hits from sequenced *Staphylococcus* and *Streptococcus* strains are retrieved from KEGG database using the SraP protein, an experimentally verified SecA2-dependent substrate (and it is the only protein that was found to be SecA2 dependent), of *Staphylococcus aureus* N315 [29]. We then used SignalP [46] to predict if these proteins have potential signal peptide, using predicted nonsecreted proteins as input for SecretomeP [48]. The final set that is predicted to be non-secretory was further analyzed by TatP [47], which can predict if these proteins can be secreted through the Twin-arginine secretion pathway. We also analyzed the original dataset by TatP to see if they had conflicting results with those predicted by SignalP. The length of truncated sequences for prediction was set to 200, as indicated previously [49].

Predicted Sec and Tat substrates were grouped according to their Cluster of Orthologous Genes (COG) functional categories. We also tested COG categories for enrichment for Sec-dependent secreted substrates using *χ*^2^:

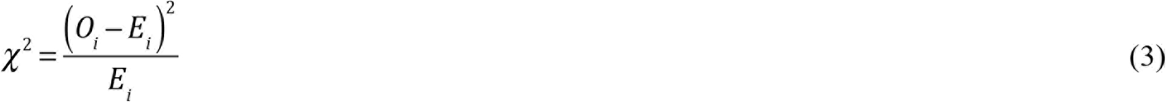

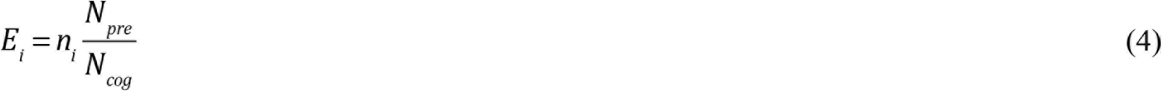

*O*_*i*_ is the observed number of predicted substrate proteins in COG category *i*. while *E*_*i*_ is the expected number of such proteins in that category *i*. The total number of proteins in category *i* is indicated by *n*_*i*_. *N*_*pre*_ is the total number of predicted substrate proteins, while *N*_*cog*_ is the total number of proteins in the proteome. Only proteins assigned within single COG functional categories were analyzed, where ambiguous categories R and S were not included.

### Intermolecular coevolutionary analysis

To understand the evolutionary relationships between domains within the same or different proteins, we tested dependencies between the evolutionary patterns of amino acid sites. We define here coevolution as the correlated variation of two amino acids, which may result from mutations in one site imposing a constraint over the mutations in another. To distinguish this type of coevolution from historical and stochastic covariation, we used a well-tested method [50] implemented in the program CAPS [51]. CAPS outperforms other methods in predicting molecular coevolution [52].

CAPS calculates the variance in the strength of mutations in an amino acid site (using BLOSUM) of a protein alignment. After correcting this variance by protein divergence levels, it is used to calculate the correlation between every two amino acid columns in the sequence alignment. These correlations are then tested for significance based on a distribution of simulated sequences (100 million pairs of sites in this study) that follow the same evolutionary dynamics as the real sequence alignments. Only correlations that are significant at a 95% level (P < 0.05), after multiple-test correction, are taken as evidence of coevolution between two amino acid sites.

Normalisation of transition scores and the correction method used in CAPS perform well when comparing sequences from different species as well as when the analyzed data are polymorphic protein sequences (allelic proteins) from the same species (e.g. human immunodeficiency virus env gene protein products gp120 and gp41 [53], human leukocyte antigen molecules [54]). Pairs of amino acid sites identified as coevolving were classed within the same group when such pairs were inclusive (for example, if site “A” was coevolving with “B”, “B” with “C” and “A” with “C”, then a group was formed including all three amino acid sites). When protein structures are available, we can identify structurally and/or functionally important coevolving sites [55]. To ensure convergence in the results, we performed the analyses three times using the same set of CAPS parameters. All three results were consistent between the three runs and reproducible in all the sequence alignments used in this study.

### Structural analysis of SecYEG translocon and SecA ATPase

To define protein domains of SecA, *E.coli* SecA domain regions [1,56] were mapped to SecA alignment. Domain information obtained here was also used for other purposes, such as, mapping sites under FD and coevolving sites to protein domains. To define protein domains of SecY, the transmembrane domain (T1-T10), cytoplasmic domain (C1-C6) and periplasmic domains (P1-P5) were obtained using SMART (Simple Modular Architecture Research Tool) service [57,58] with *E.coli* SecY protein sequence.

To understand the structural and functional interactions between amino acid sites under FD and coevolution, protein X-ray crystallographic complete structures for SecA, SecYEG-SecA complex were retrieved from Protein Data Bank (PDB, http://www.pdb.org). The PDB codes of these structures are 1M6N [11], 1NKT [56], 1TF2 [59], 2IBM [60], 2IPC [61], 3JUX [62], 3JV2 [62] and 3DIN [12]. SecA from SecYEG-SecA (3DIN, Figure 3H) was used as the comparison base when performing the structure based alignment through CEalign in PyMOL (Figure 3A-H). PyMOL Version 1.3 (http://pymol.sourceforge.net/) was used to visualize X-ray crystallographic structure, and the structure alignment was performed using CEalign plug-in (http://pymolwiki.org/index.php/Cealign) in PyMOL [63]. We calculated the spatial distance between two amino acids by taking their shortest atomic Euclidean distance. We followed the same procedure for every pair of atoms in both proteins, and then we took the shortest atomic distance as the amino acid distance between both of the amino acids. We followed the same procedure as well to calculate the shortest amino acid atomic distances between domains and between amino acid sites under FD and those identified in [64] as drug targets.

## Results

We screened a large bacterial phylogeny to identify clades with strong evidence of molecular adaptive changes, indicative of functional divergence.

### Evidence of functional divergence of the motor and translocon pore in pathogenic bacteria

We identified 298, 87, 191, 199 and 268 clades that have at least one amino acid site under FD (Table S1 to S5 for SecA, SecB, SecE, SecG and SecY, respectively). We found 271 (35% of all species under FD), 89 (31% of all species under FD), 189 (29% of all species under FD), 199 (30% of all species under FD) and 219 (29% of all species under FD) pathogenic bacterial strains to have at least one amino acid site under FD in SecA, SecB, SecE, SecG and SecY, respectively. Interestingly, the clades with the strongest signal of FD included clinically important pathogenic bacteria (Table S1 to S5), such as *Chlamydia*, *Listeria, Staphylococcus* and *Streptococcus* (Figure 4).

Enrichment analyses of FD (Figure 1A and Figure 2A) showed that the domain most affected by FD was NBD1, involved in binding and hydrolysis of ATP. Also, NBD1 and PPXD domains present similar FD heatmap patterns compared to other domains. HSD and NBD2 domains of SecA were clustered within the same FD enrichment group. CTL and HWD domains were clustered as being the most impoverished for FD (Figure 1A), indicating that their functions have been conserved during SecA evolution. Taking all clades into account instead of the 10% most enriched ones made no difference to our results (Figure S1).

**Figure 2.**
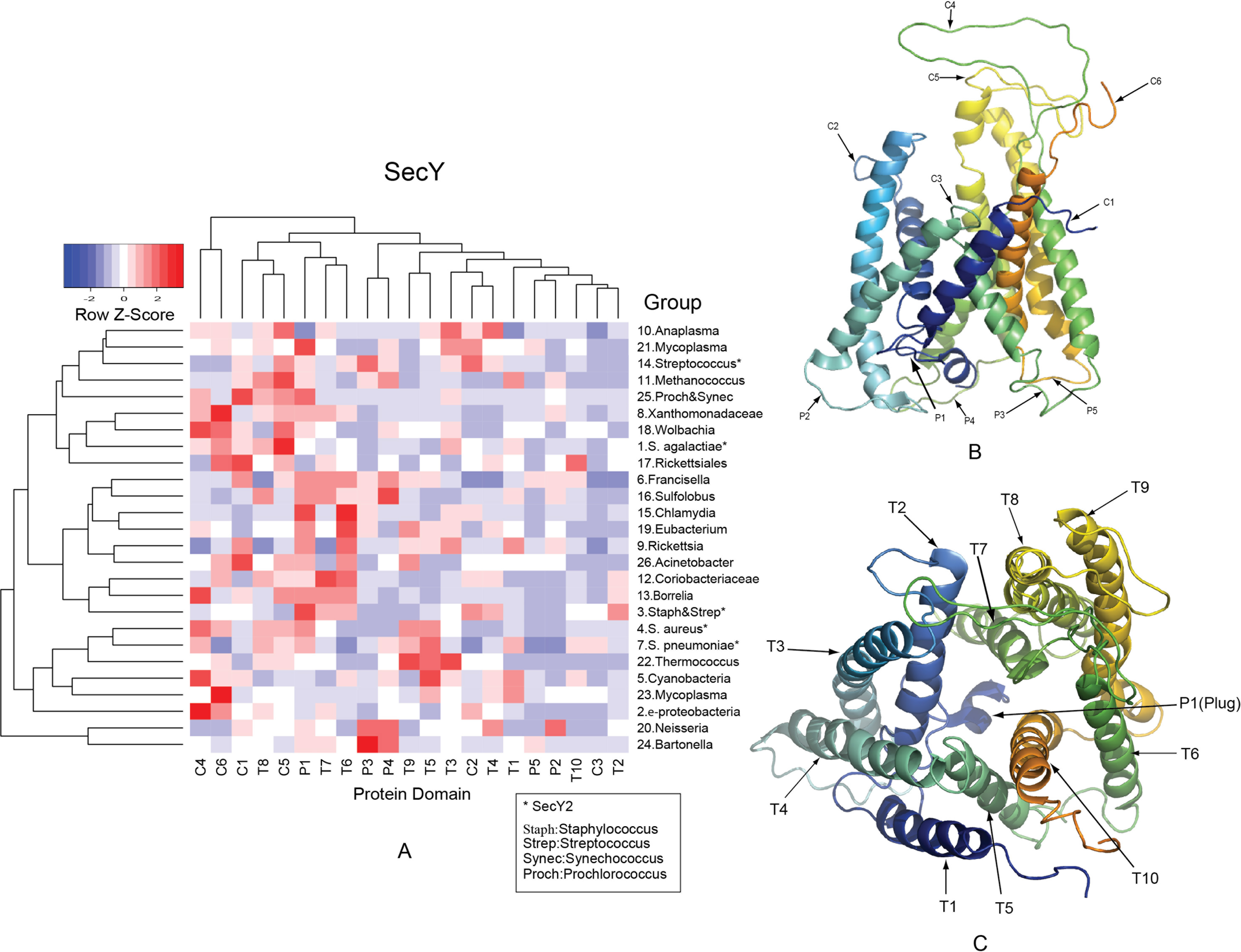
Two-way clustering of the top 10% clades under functional divergence in SecY. (A) Protein domains are clustered according to the number of sites under functional divergence after Z-score standardization, group number represents the rank of number of sites under functional divergence in each analyzed branch, and groups are also clustered. (B) Cytoplasmic (C1-C6) and periplasmic (P1-P5) domains are shown in SecY. (C) Transmembrane (T1-T10) domains are shown in SecY. The so-called “plug” domain is also shown (P1) in the centre. * denotes functional divergence results for SecY2, representative phylogenetic group in each group is also shown.

At the structural level, amino acids under FD are not clustered in a single region but present significant distances between one another in all main conformational SecA states (table 1, and Figures 3A to 3H). In particular, amino acid sites under FD between the PPXD domains and the remaining domains (NBD2, NBD1 and HWD) deviates the most in terms of mean atomic distance, especially to the sites from the NBD2.

**Figure 3.**
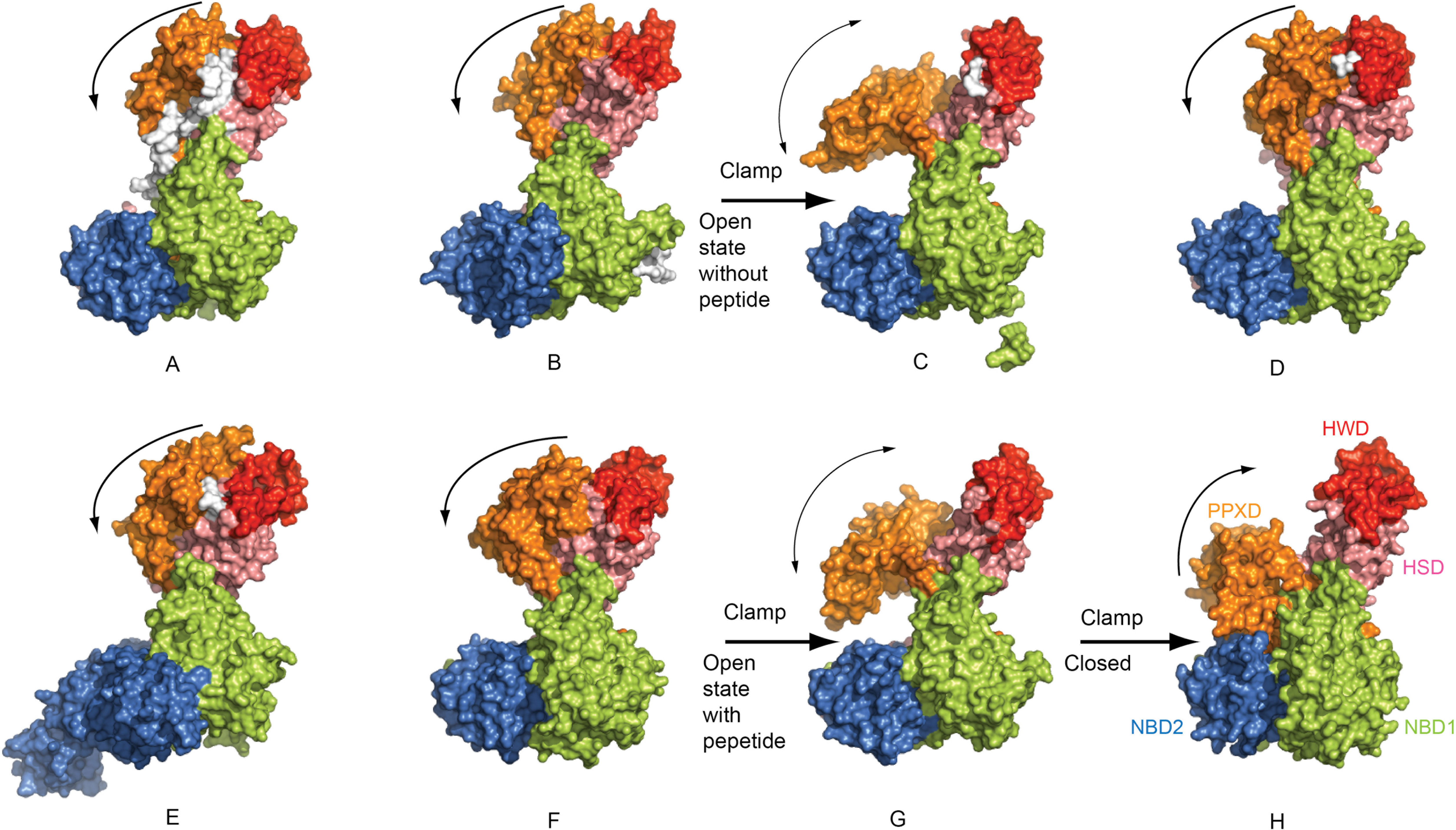
Structural alignment of SecA. Solved SecA crystal structures are aligned based on the structure for (H). The structures are presented for (A) 1M6N [11], (B) 1NKT [56], (C) 1TF2 [59], (D) 2IBM [60], (E) 2IPC [61], (F) 3JUX [62], (G) 3JV2 [62]. The possible trend of domain movement of PPXD is denoted by arch lines with arrow.

**Figure 4.**
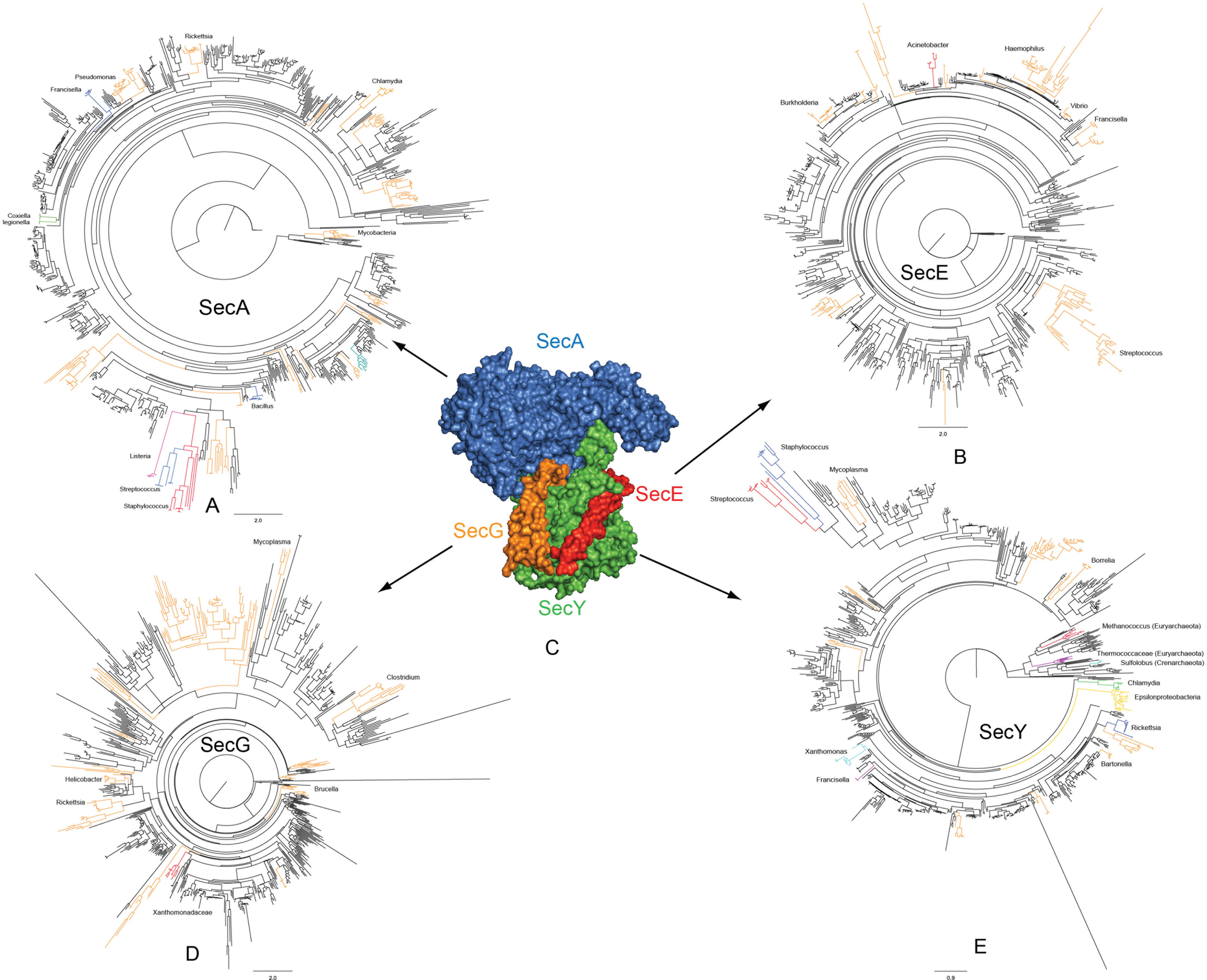
Protein components SecA (A), SecE (B), SecG (D) and SecY (E) under functional divergence of the SecA-SecYEG complex (C) and their corresponding phylogenetic trees, where the top 10% most functionally divergent clades (see Table S1, S2, S3, S4 and S5 for details) are color labeled on their corresponding phylogenetic trees.

**Table 1.** Mean domain distance of amino acid sites under functional divergence. The mean distances between amino acid sites under functional divergence from different protein domains (NBD1, PPXD, NBD2, HSD and HWD) of SecA with four different functional states are calculated. PDB IDs of SecA structures are IM6N, 1TF2, 3JV2 and 3DIN, and corresponding SecA structures are shown in Figure 4.

| Domain1 | Domain2 | Mean Amino Acid Distance ( Å) |  |  |  |
| --- | --- | --- | --- | --- | --- |
|  |  | IM6N (Figure 3A) | 1TF2 (Figure 3C) | 3JV2 (Figure 3G) | 3DIN (Figure 3H) |
| NBD2 | NBD1 | 37.78 | 37.21 | 37.1 | 37.19 |
| NBD2 | HWD | 75.32 | 74.52 | 74.25 | 74.22 |
| NBD2 | PPXD | 54.99 | 42.21 | 38.98 | 31.18 |
| NBD2 | HSD | 48.82 | 47.94 | 48.08 | 46.3 |
| NBD1 | HWD | 56.3 | 57.5 | 56.8 | 63.69 |
| NBD1 | PPXD | 49.11 | 46.11 | 44.44 | 40.44 |
| NBD1 | HSD | 41.11 | 41.58 | 41.6 | 44.48 |
| HWD | PPXD | 29.66 | 43.96 | 45.35 | 47.87 |
| HWD | HSD | 28.97 | 29.73 | 29.35 | 31.42 |
| PPXD | HSD | 25.42 | 29.3 | 28.8 | 29.17 |

In SecY (SecY1 and SecY2) domains C4 and C6 cluster together in terms of enrichment for FD (Figure 2A). Previous studies highlighted an important role of C4 and C6 domains in SecY function [13,66] and their FD may have important functional consequences in the performance of SecY. Like in SecA, SecY showed similar clustering patterns when the full set of bacterial clades was considered (Figure S2). Domains C1, T8, C5, P1, T7 and T6 form the second cluster, and P3, P4, T9, T5, T3, C2, T4, T1, P5, P2, T10, C3 and T2 form the third cluster. This clustering is supported by some functional data. For example, T8, T7 and T6 are involved in the plug domain P1 displacement for subsequent peptide translocation upon ribosome [67] or SecA [12] binding to C1 (C-terminal tail) and C5 domains of SecY.

### Functional divergence of pathogenic SecA1/SecY1

How did the function of the canonical SecA/SecY diverged across the bacterial phylogeny? We found that SecA and SecY presented very different FD patterns across the bacterial phylogeny (Figures 1A and 2A). The clustering of non-sister bacterial clades within the same FD-enrichment group indicates, nevertheless, that unrelated bacterial groups have undergone similar FD events, allowing them colonise similar ecological niches. In concert with this prediction, pathogenic bacteria, such as *Chlamydia* (group 1, Figure 1A and 4D), presented strong evidence for FD. *Chlamydia* also presented a unique FD profile, mainly affecting domains PPXD and NBD1, and much less HSD and NBD2. The fifth most functionally divergent group comprises five strains of *Coxiella burnetii*, clustered with group 7 (group 5 and 7 for SecA, Figure 1A) including *Mesoplasma florum, Mycoplasma capricolum* and *Mycoplasma mycoides*.

We identified 40 sites in SecY under FD between ɛ-proteobacteria (group 2 in Figure 2A) and other proteobacteria groups (α, β, γ, δ), 11 sites between α-, β-, γ- (group 102 in Table S4) and δ-proteobacteria groups, eight sites between α- (group 101 in Table S4) and β- and γ- proteobacteria groups, and six sites between β- (group 185 in Table S4) and γ- proteobacteria group (Figure 5). Sequenced bacteria from ε-proteobacterial group represent the second most functionally divergent group for SecY protein compared with other major sequenced proteobacteria (α, β, γ, δ) groups. ε-proteobacterial (group 2) showed a distinctive molecular pattern of domain FD compared to other clades (Figure 2A). To understand the relative functional importance of this pattern, we mapped sites under FD of ε-proteobacterial to the C4 loop formed between T7 and T8 (Figure 6). The tip of C4 domain is embedded in the “clamp” formed by PPXD, NBD2, NBD1 and part of the HSD (Figure 6A). 250Gly, 243Gln and 248Val have been identified under FD in this bacterial group and, given their position in C4 (Figure 6B), they are likely to interact with pre-proteins and participate in their translocation. Also, residues 255Gln, 256Gly and 257Ala of SecY interact with PPXD domain (Figure 6C and 6D) and may contribute to the movement of C4 loop.

**Figure 5.**
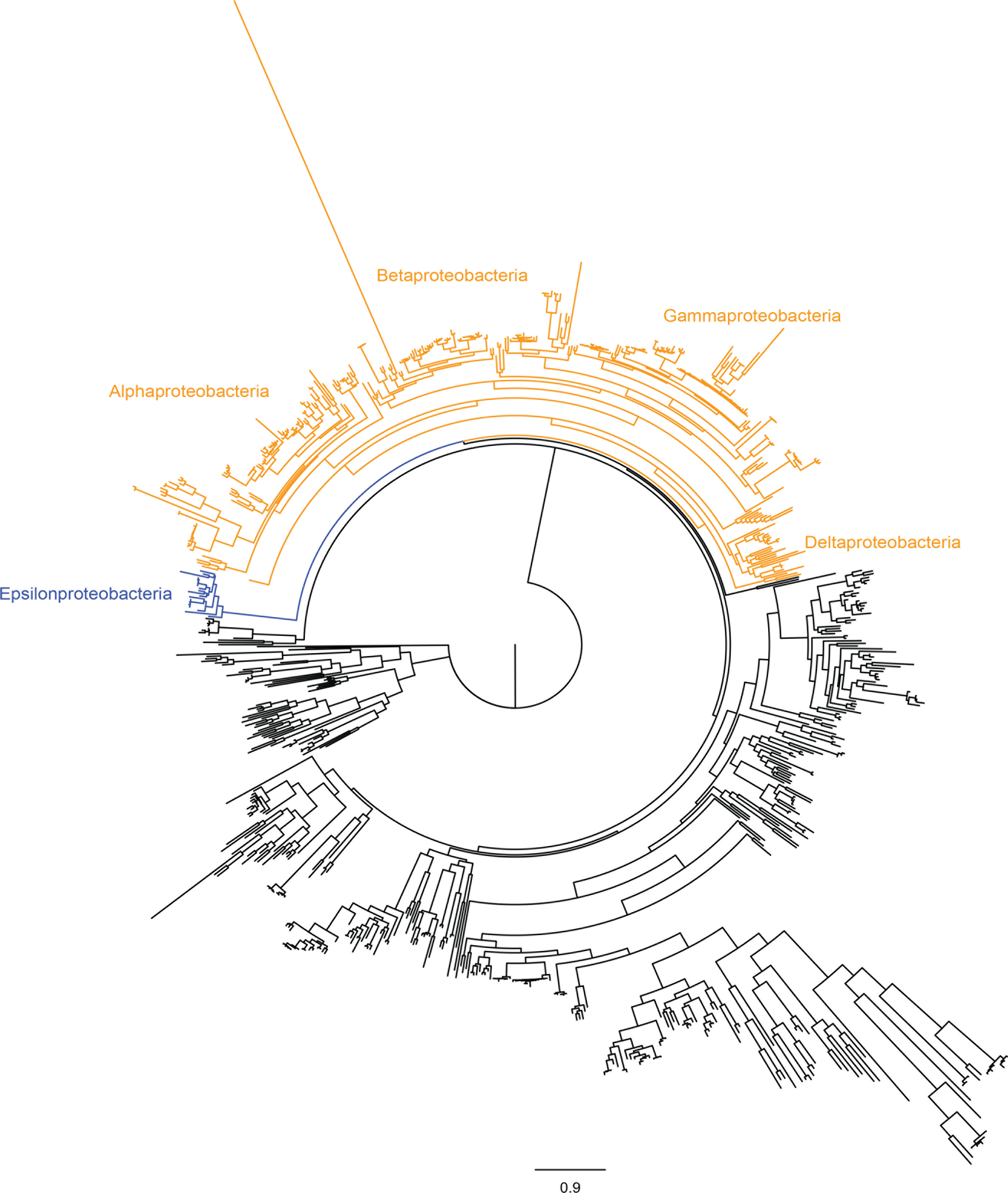
Maximum likelihood phylogenetic tree of SecY. Proteobacteria groups are mapped to the tree and the different groups are identified, being these epsilonproteobacteria, alphaproteobacteria, betaproteobacteriam gammaproteobacteria and deltaproteobacteria.

**Figure 6.**
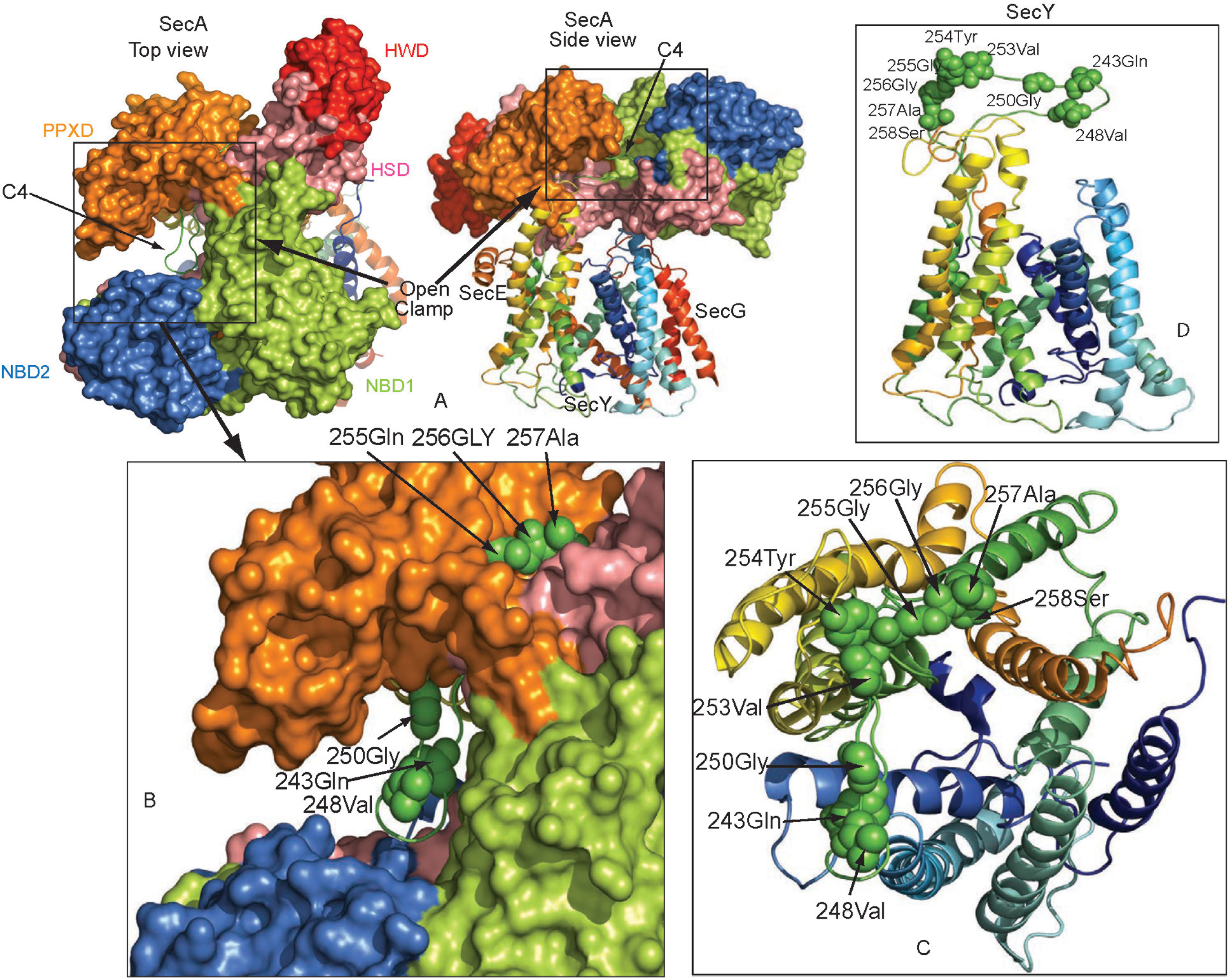
Amino acid sites mapped to the C4 domain of SecY. Amino acid sites under functional divergence from s-proteobacteria were mapped to the three-dimensional crystal structure of SecYEG-SecA complex. (A) Three-dimensional crystal structure of SecA from *B. subtilis* (PDB: 3JV2) was superimposed to the SecYEG-SecA structure using CEalign in PyMOL. The original SecA structure was then removed from the SecYEG-SecA complex. We show a top view of the complex in the left side figure of (A) and a side view of this complex in the right side of (A). (B) A zoomed in figure of the C4 region clearly showed that mapped amino acid residues are in the clamp and also interact with the PPXD domain. (C) The same figure as shown in (B) but excluding SecA region, in which all sites from the C4 domain can be clearly observed from the top of SecY structure. (D) The same sites are viewed from the SecY side.

### Functional specialization of SecA2/ SecY2

FD in Sec system may have been required in some pathogenic bacteria to secrete large serine-rich glycoproteins. These proteins include SraP and its homologs in *Staphylococcus* and *Streptococcus* species [29] (Table 2). A recent study has shown that both SecA1 and SecA2 proteins are essential in *Corynebacterium glutamicum* [68]. In contrast with the canonical Sec system, here we found that all SecA2 showed greater FD compared to SecA1 in those bacteria that have both SecA1 and SecA2 proteins, with the only exception being *Corynebacterium spp*. (group 14, Figure 1A). The SecA2/Y2 system of *Streptococcus spp*. and *Staphylococcus spp*. (group numbers 10, 11, 16, 19 and 1, 3, 4, 7, 14 for SecA2 (Figure 1A, Figure 4A) and SecY2 (Figure 2A, Figure 4E), respectively) are among the top 10% most functionally divergent bacterial clades.

**Table 2.** Orthologs of *Staphylococcus aureus* N315 SraP from Staphylococcus and Streptococcus species.

| Strains | Gene Locus | Cleavage Site | Prediction Method |
| --- | --- | --- | --- |
| *Staphylococcus aureus* N315 | SA2447 (SraP) | 90 and 91 | SignalP |
| *Staphylococcus aureus* COL | SACOL2676 | 90 and 91 | SignalP |
| *Staphylococcus aureus* ED98 | SAAV_2725 | 90 and 91 | SignalP |
| *Staphylococcus aureus* JH1 | SaurJH1_2734 | 90 and 91 | SignalP |
| *Staphylococcus aureus* JH9 | SaurJH9_2678 | 90 and 91 | SignalP |
| *Staphylococcus aureus* MSSA476 | SAS2540 | NA | SecretomeP |
| *Staphylococcus aureus* Mu3 | SAHV_2638 | 90 and 91 | SignalP |
| *Staphylococcus aureus* Mu50 | SAV2654 | 90 and 91 | SignalP |
| *Staphylococcus aureus* MW2 | MW2575 | NA | SecretomeP |
| *Staphylococcus aureus* NCTC8325 | SAOUHSC_02990 | 90 and 91 | SignalP |
| *Staphylococcus aureus* Newman | NWMN_2553 | 90 and 91 | SignalP |
| *Staphylococcus aureus* USA300 | SAUSA300_2589 | 90 and 91 | SignalP |
| *Staphylococcus aureus* USA300 TCH1516 | USA300HOU_2654 | 90 and 91 | SignalP |
| *Staphylococcus carnosus* | Sca_2202 | 32 and 33 | SignalP |
| *Staphylococcus epidermidis* ATCC 12228 | SE2249 | NA | SecretomeP |
| *Staphylococcus epidermidis* RP62A | SERP2281 | NA | SecretomeP |
| *Staphylococcus haemolyticus* | SH0326 | 92 and 93 (17 and 18) | SignalP (TatP) |
| *Staphylococcus saprophyticus* | SSP0135 | 50 and 51 | SignalP |
| *Streptococcus agalactiae* A909 (serotype Ia) | SAK_1493 | 50 and 51 | SignalP |
| *Streptococcus agalactiae* NEM316 (serotype III) | gbs1529 | NA | SecretomeP |
| *Streptococcus gordonii* | SGO_0966* | 90 and 91 (Reference) | SecretomeP |
| *Streptococcus pneumoniae* 70585 | SP70585_1816* | NA | SecretomeP |
| *Streptococcus pneumoniae* ATCC 700669 (serotype 23F ST81 lineage) | SPN23F_17820* | NA | SecretomeP |
| *Streptococcus pneumoniae* CGSP14 (serotype 14) | SPCG_1750* | NA | SecretomeP |
| *Streptococcus pneumoniae* Hungary19A 6 | SPH_1885* | NA | SecretomeP |
| *Streptococcus pneumoniae* TIGR4 (virulent serotype 4) | SP_1772 | 42 and 43 | SignalP |
| *Streptococcus sanguinis* | SSA_0829* | NA | SecretomeP |

We identified 14, 8, 16, 14, 3 and 0 sites under FD in the NBD1, PPXD, NBD2, HSD, HWD and CTL domains of SecA, respectively (Figure 7). Moreover, domains NBD2, HSD and NBD1 were specially enriched for sites under FD (Figure 1A), and some of these sites in SecA2 are located in two regions recently identified as functionally important. First, amino acids 774Glu, 775Ala, 786Pro and 793Glu are located in the two helix finger of the HSD domain (Figure 7D) and have been suggested to play an important role in moving polypeptide chains into the SecY channel [69]. Second, amino acid site 77Phe was identified in NBD1 (Figure 7E) and has been identified as major targets for SecA ATPase inhibitor [64,70]. 35 of the sites with FD were mapped to the C1-C6, T1-T10 and P1-P5 domains of SecY (sites located on unsolved structure regions are not shown), (Figure 8). Because of their greater enrichment for FD (group 4 in Figure 2A), domains C4, C6, T8, C5, P1, T6 and T5 may play important functions in the adaptation to novel ecological niches.

**Figure 7.**
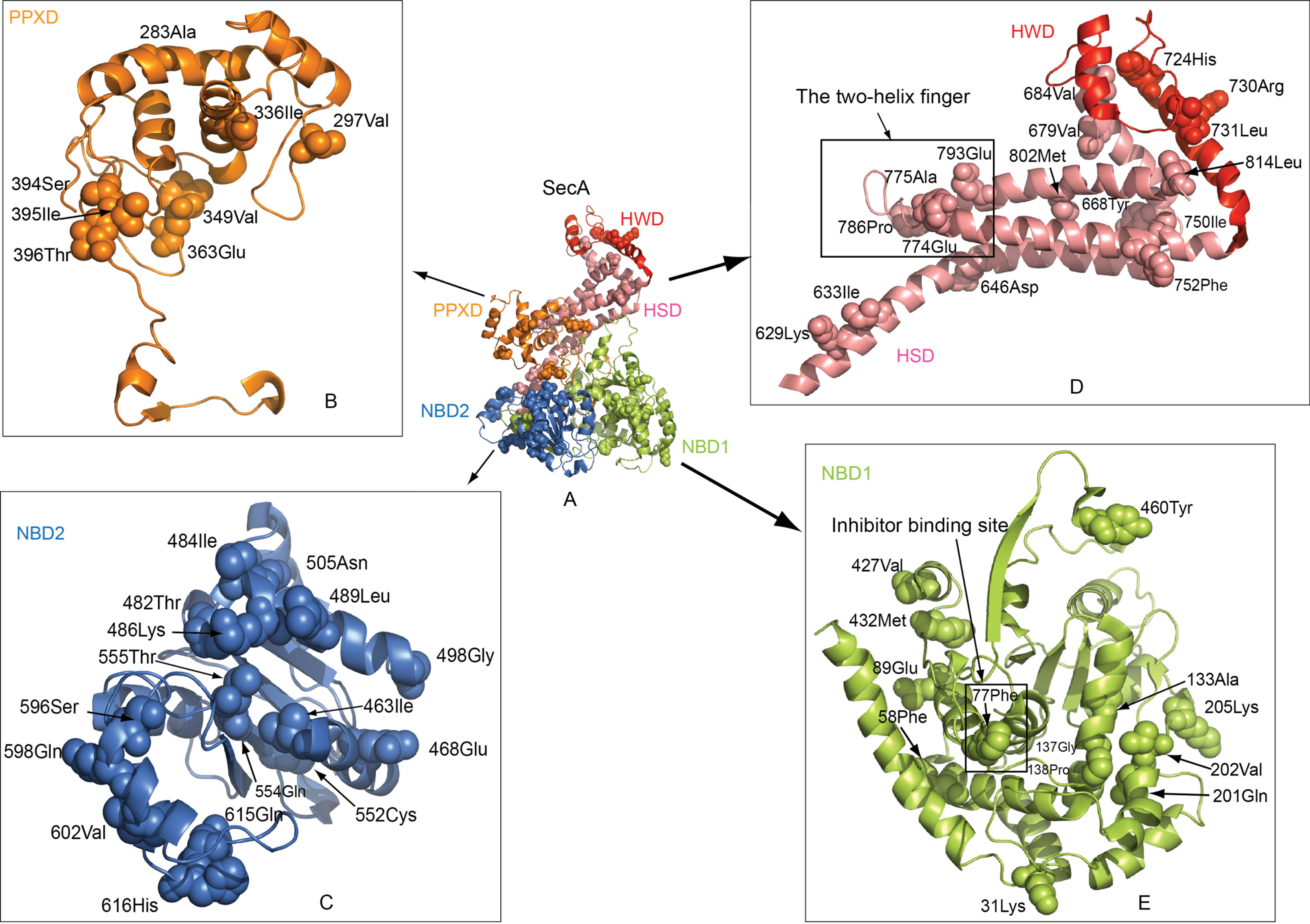
Amino acid sites under functional divergence of *Staphylococcus spp*. (group 10 in Figure 2) mapped to SecA. (A) Sites are mapped to domains PPXD, NBD2, HSD, HWD and NBD1 of SecA. (B) Detailed view of sites at PPXD. (C) Detailed view of sites at NBD2. (D) Detailed view of sites at HSD and HWD. (E) Detailed view of sites at NBD1.

**Figure 8.**
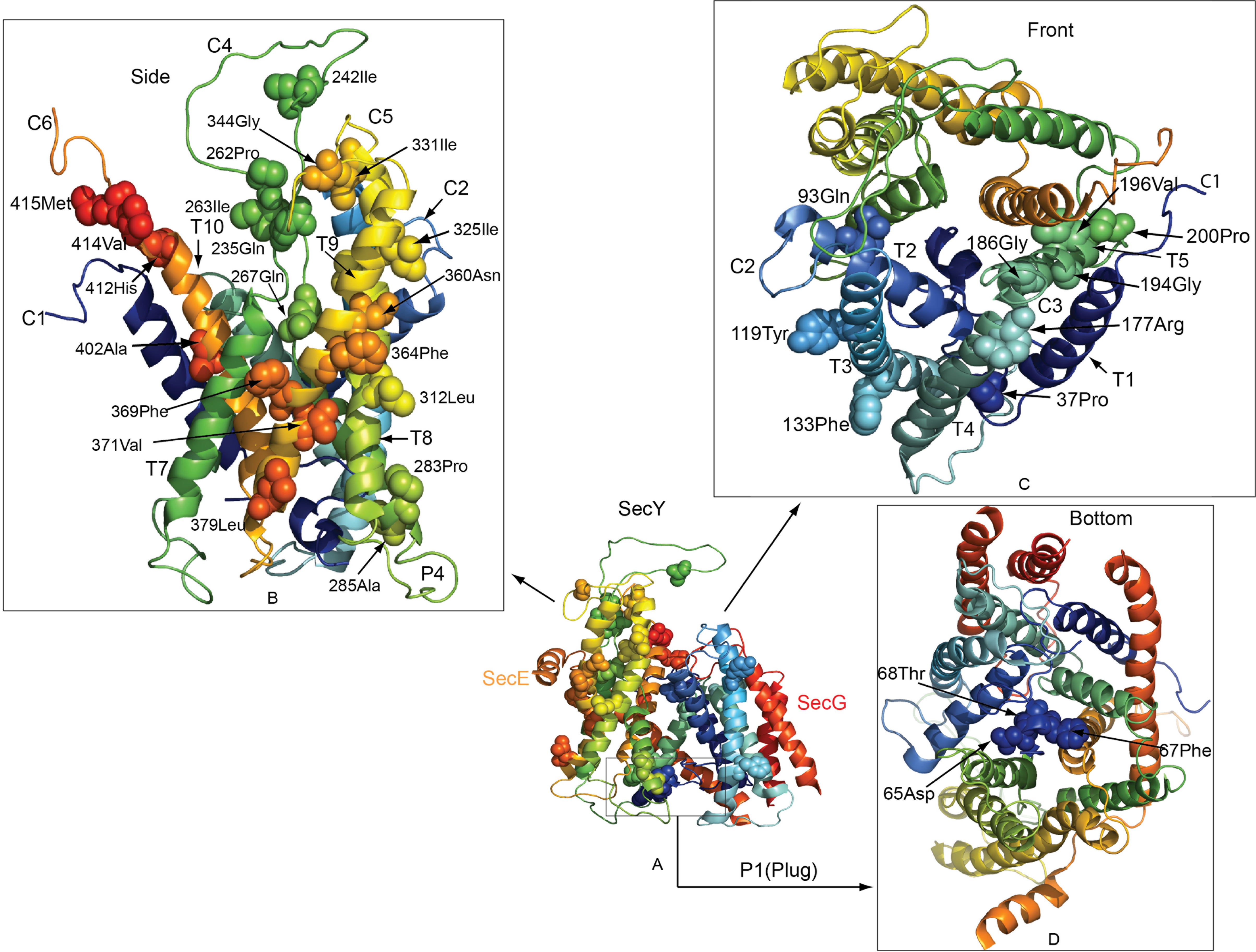
Amino acid sites under functional divergence of *Staphylococcus spp*. (group 4 in Figure 3) mapped to SecY. (A) Sites are mapped to SecYEG. (B) Detailed view of sites from SecY side. (C) Detailed view of sites from SecY front. (D) Detailed view of sites from SecY bottom.

Remarkably, among the bacteria with evidence for FD, 11 *Mycobacteria* that lack SecY2 (group number 22 in SecA2, Figure 1A) were clustered with group 28 consisting of seven plant species *(Physcomitrella patens subspp. patens, Sorghum bicolor, Vitis vinifera, Arabidopsis thaliana, Populus trichocarpa, Oryza sativa japonica* and *Ricinus communis*).

Because SecA2 and SecY2 have suffered dramatic evolutionary changes, we seeked to understand whether these two proteins maintained a reciprocal evolutionary dependency. To this end we applied a method to identify molecular coevolution. Coevolution analysis identified two groups of residues that involved amino acids from SecA2 and SecY2. Group 1 has 8 coevolving residues (Figure 9A and Table 3), while group 2 has 19 co-evolving residues (Figure 9B, Table 3). The two groups of coevolution include residues from T3, T8 and T10 domains of SecY2 (Table 3). Importantly, these domains have been shown to play an important role in the interchange between the open and close states of the SecY channel and in the displacement of the channel plug (P1 region). Co-evolving residues (465Glu and 593Ile) and 44Trp in group 2 are located in the NBD2 domain of the motor protein SecA2 and P1 domain (the plug domain) of SecY2 (Figure 9B). The plug domain is believed to play a role in sealing and opening the translocation channel SecY [12,71–78]. Moreover, an amino acid site (396Thr) located in the PPXD domain was found to strongly coevolve with a high number of amino acids from SecY2 (326 amino acid sites), indicating an important role of this site from the PPXD domain in translocating SecA2-dependent substrate proteins.

**Figure 9.**
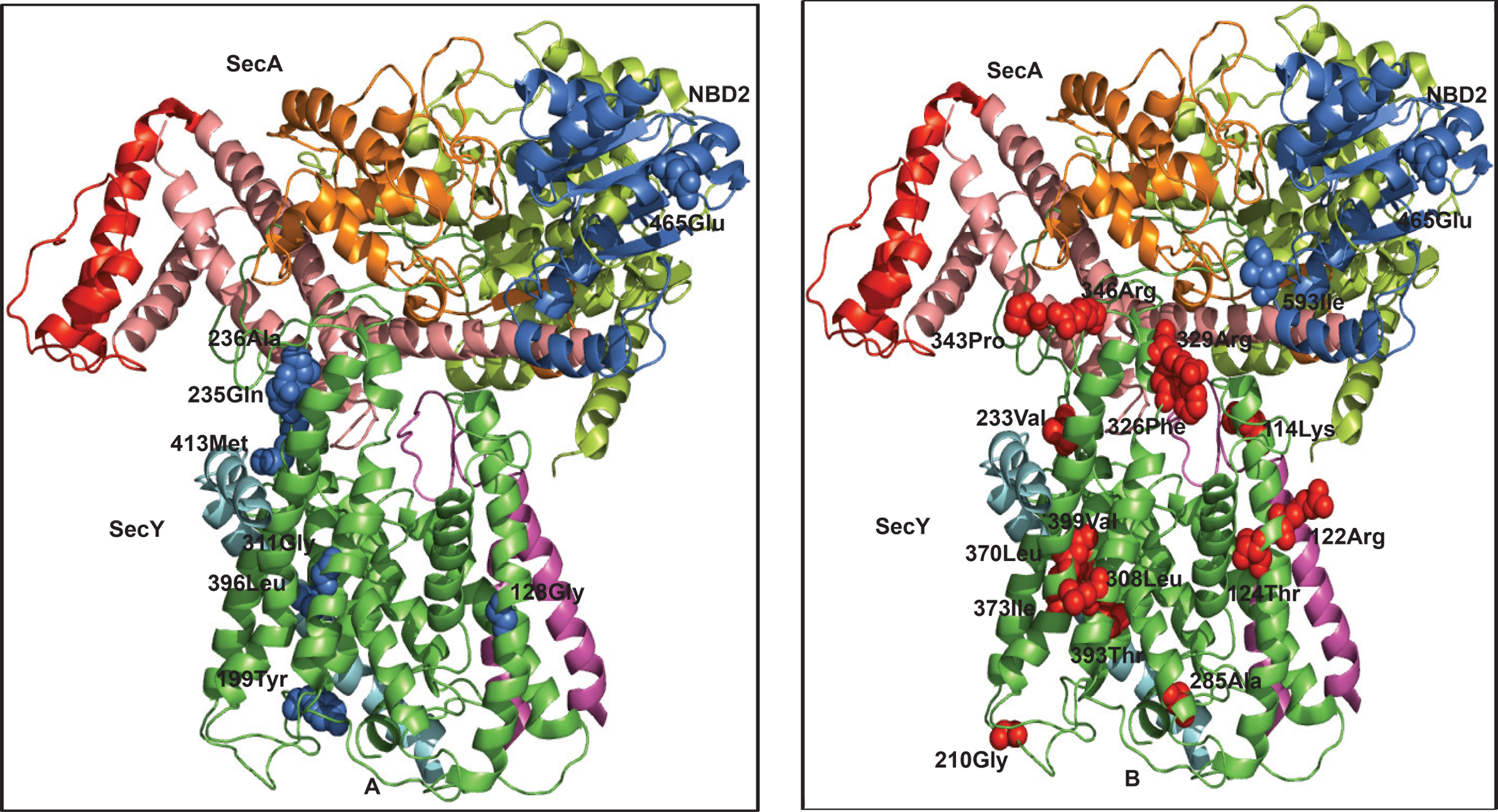
Coevolving residues of the accessory SecA2/SecY2 system mapped to SecYEG- SecA complex. (A) Coevolving amino acid residues of group one (Table 3) are mapped to SecA (blue balls) and SecY (blue balls), respectively. (B) Coevolving amino acid residues of group two (Table 3) are mapped to SecA (blue balls) and SecY (red balls), respectively.

**Table 3.** Functionally coevolving amino acid sites from the accessory SecA2/SecY2 system. Sites are mapped to domains and proteins and classified into two groups by CAPS analysis.

| Group | AA Site | Domain | Protein |
| --- | --- | --- | --- |
| 1&2 | 465Glu | NBD2 | SecA2 |
| 2 | 593Ile | NBD2 | SecA2 |
| 1 | 236Ala | C4 | SecY2 |
| 1 | 235Gln | C4 | SecY2 |
| 1 | 413Met | C6 | SecY2 |
| 1 | 128Gly | T3 | SecY2 |
| 1 | 199Tyr | T5 | SecY2 |
| 1 | 311Gly | T8 | SecY2 |
| 1 | 396Leu | T10 | SecY2 |
| 2 | 4Ala* | C1 | SecY2 |
| 2 | 114Lys | C2 | SecY2 |
| 2 | 346Arg | C5 | SecY2 |
| 2 | 329Arg | C5 | SecY2 |
| 2 | 343Pro | C5 | SecY2 |
| 2 | 44Leu* | P1(Plug) | SecY2 |
| 2 | 210Gly | P3 | SecY2 |
| 2 | 393Thr | T10 | SecY2 |
| 2 | 399Val | T10 | SecY2 |
| 2 | 124Thr | T3 | SecY2 |
| 2 | 122Arg | T3 | SecY2 |
| 2 | 233Val | T6 | SecY2 |
| 2 | 285Ala | T7 | SecY2 |
| 2 | 326Phe | T8 | SecY2 |
| 2 | 308Leu | T8 | SecY2 |
| 2 | 373Ile | T9 | SecY2 |
| 2 | 370Leu | T9 | SecY2 |

### Functional divergence in SecB

We identified a clade with the strongest FD (including 20 amino acids under FD) to contain mostly pathogenic *Rickettsia spp*. The high similarity of SecB structures [79] to that of *Haemophilus influenzae* (root mean square deviation: RMSD = 3.2 angstroms), indicates that the over-all protein structure of SecB is highly conserved among bacteria. Most FD sites were distributed between two SecB regions, which are homologous to binding-regions in SecA (Figure 10). One is close to the C-terminal linker on SecA [80] (Figure 10A). The other is close to the SecB-binding site on SecA (Figure 10B). This SecB-binding site is highly conserved and located in the C-terminus of SecA, hence functionally divergent amino acid sites on SecB may play a role in regulating SecA by binding to the conserved C terminus region of this protein. This is consistent with our FD analysis of SecA, where we show that the CTL domain is impoverished for FD (Figure 1A), suggesting that the possible ATPase activity of CTL domain in SecA is highly conserved among bacteria groups.

**Figure 10.**
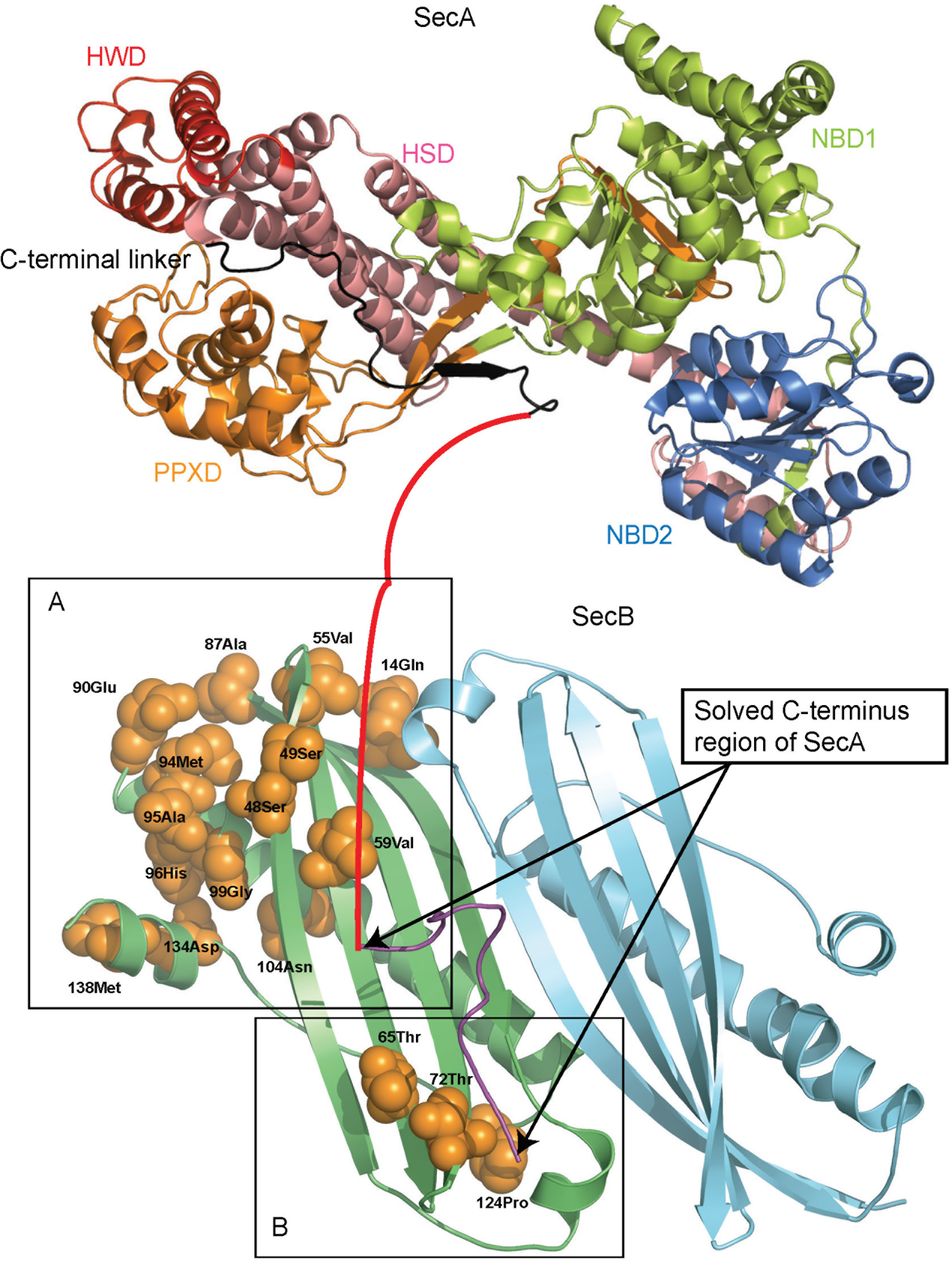
Amino acid sites under functional divergence in *Rickettsia spp*, (group 1 in Table S2) mapped to SecB structure. SecA structure is shown to demonstrate the possible interactions between SecA CTL domain and SecB dimer (left one is colored green, right one is colored cyan, sites are only mapped to the left one). (A) SecA-binding site close to C-terminal linker (colored black on the SecA structure), note that unsolved CTL region is colored red, it is manually linked to the solved C terminus region (for illustration purpose only). (B) SecA-binding site close to the highly conserved C terminus region (colored magenta, solved structure from PDB: 1QYN, 1OZB).

To address the evolutionary relationship between CTL and other protein domains in SecA, SecB and SecY, we performed coevolutionary analyses between SecA and SecY and SecA and SecB, respectively. In the combined coevolutionary networks between SecA, SecB and SecY, we found that three amino acid sites in the CTL domain from SecA were responsible for the significant coevolutionary patterns found between the three proteins. Amino acids coevolving with these three belonged to most of the domains in SecY (except C2 and T2) and were also identified in the regions in SecB homologous to the two binding regions of SecA (Figure 10A and Figure 10B). These results suggest that CTL domain may regulate other protein domains during the interaction of SecB, SecA and SecY (Supplementary Figure S3).

### Functional divergence in SecE and SecG

A clade containing two pathogenic *Acinetobacter spp*. was the most functionally divergent group of SecE (Figure 4B). Interestingly, we observed that SecG proteins of plant pathogens from *Xanthomonas spp*. and *Xylella fastidiosa* have accumulated the most radical changes (Figure 4C) indicating that SecG may play a role in the pathogenesis of bacteria in plants. We mapped the sites under FD to SecE (Figure 11A) and SecG (Figure 11B) in the SecYEG-SecA structure. In SecE, sites 49Phe, 32Val and 24Lys are in the vicinity of the transmembrane domains of SecY, and they may consequently contribute to the proper function of SecY channel. Similarly, in SecG, sites 29Glu, 26Lys, 65Val, 66Ser and 73Val may contribute to the proper function of SecY and possible interaction with SecA.

**Figure 11.**
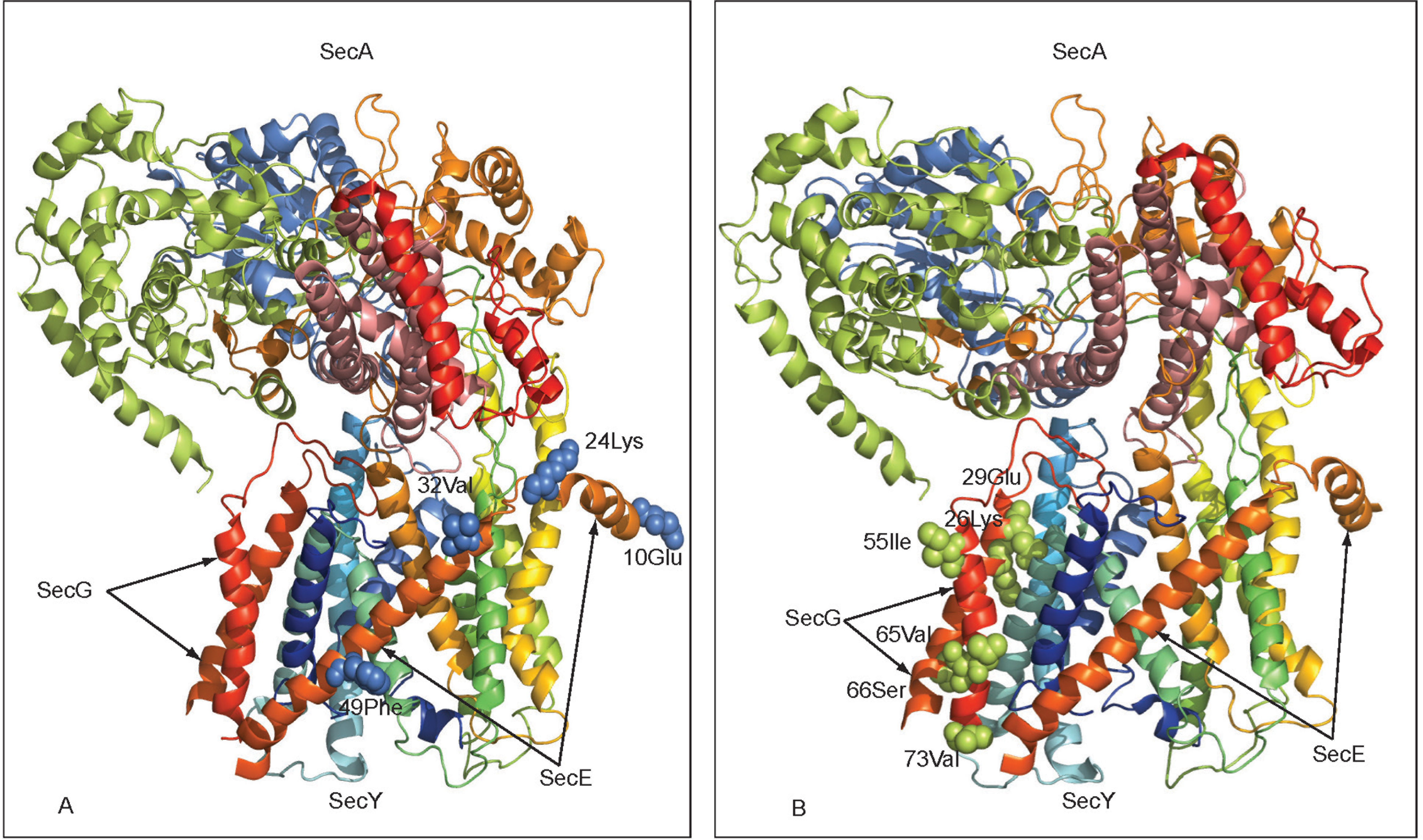
Amino acid sites under functional divergence (group 1 in Table S3 and Table S4) of SecE and SecG mapped to the SecA-SecYEG structure. (A)Sites are mapped to SecE. (B) Sites are mapped to SecG.

### Functional divergence of translocon core protein SecY in archaeal groups

There are 24 archaeal clades identified with at least one site under FD (Table S4). Three archaeal groups (group number 11, 16 and 22, Figure 2A) were among the top 10% most functionally divergent: *Methanococcus* and *Methanocaldococcus spp*. (group number 11), *Sulfolobus spp*, (group number 16), *Thermococcus spp*. and *Pyrococcus spp*. (group number 22). Interestingly, group 11 and 22 were not clustered with any other bacterial groups (Figure 2A), indicating a unique pattern of FD. *Thermococcus spp*, and *Pyrococcus spp*, are hyperthermophilic archaea and they belong to the order *Thermococcales*. *Thermococcales* can be found in terrestrial, submarine hot vents and deep subsurface environments. Importantly *Pyrococcus spp*. can only be found in marine environments and belongs to a particular niche [81]. It is tempting to speculate that the above clustering pattern for group 11 and 22 may be a consequence of adaptation to their specific living environments.

### Differential protein secretion in functionally divergent bacteria

To find possible correlations between functional divergence and ecological adaptation, we identified the Sec- and Tat-dependent secreted proteins of the top 5 most functionally divergent clades for SecA, SecB, SecE, SecG and SecY, respectively (we analyzed 207 secretomes, for complete results see Supplementary Table S6). To characterize the functions of these secreted proteins, we grouped these proteins into COG functional categories and statistically tested their significance. We considered a group to be enriched or impoverished for secreted proteins if *P* <0.01. First, most secretomes were enriched in COG category M (Cell wall/membrane/envelope biogenesis). This is expected, as the Sec system is a major secretion pathway for secreting membrane related proteins. Second, we unexpectedly found that COG category P (Inorganic ion transport and metabolism) was enriched in almost all the secretomes of pathogenic as well as non-pathogenic bacteria under functional divergence. This indicates that proteins responsible for inorganic ion transport and metabolism may play an important role in bacterial niche adaptation. This observation however would require experimental testing for verification. Finally, COG category U (Intracellular trafficking, secretion, and vesicular transport) was enriched in most pathogenic bacteria.

Analysis of enrichment for the 5 representative species from clades that have the largest number of sites under FD for Sec proteins show that obligate intracellular pathogen *Rickettsia prowazekii* has COG category O (Posttranslational modification, protein turnover, chaperones) enriched in its secretome (Figure 12B, *P* < 0.01). Chaperones are known to buffer the deleterious effects of mutations by folding proteins into their functional conformation despite destabilizing mutations. Important functional mutations at chaperones may improve their folding function helping bacteria, such as *Rickettsia prowazekii*, adapt to an obligate intracellular life style. Three *Streptococcus spp*. are enriched in COG category M, P, T and U (Figure 13). Strikingly, all three *Staphylococcus* species are enriched in COG category P (P<0.001) and they had the highest number of secreted proteins in category P compared to other categories. How do secreted proteins that contribute to inorganic ion transport and metabolism contribute to *Staphylococcus spp*. niche adaptation? *Staphylococcus* species were found to survive in high salt concentrations, and it was reported that these species can grow well at NaCl concentrations as high as 10-15% or even higher [83]. To survive high salt concentrations, the cell must be able to counterbalance the osmotic pressure produced by this environment. Inorganic ions such as *Na*^+^ and *K*^+^ must be processed to maintain appropriate osmotic pressure in the cell. It is also known that pathogenic bacteria use inorganic ion concentrations to sense their location. This is important for *Streptococcus* and *Staphylococcus* species because they are frequently found to be part of the microbiota of the human skin, mucosal surface and so on [84,85]. More importantly, many crucial ions that pathogenic bacteria require are located in the host cell and bound to host proteins. Therefore, these pathogens need to actively acquire these ions [86].

**Figure 12.**
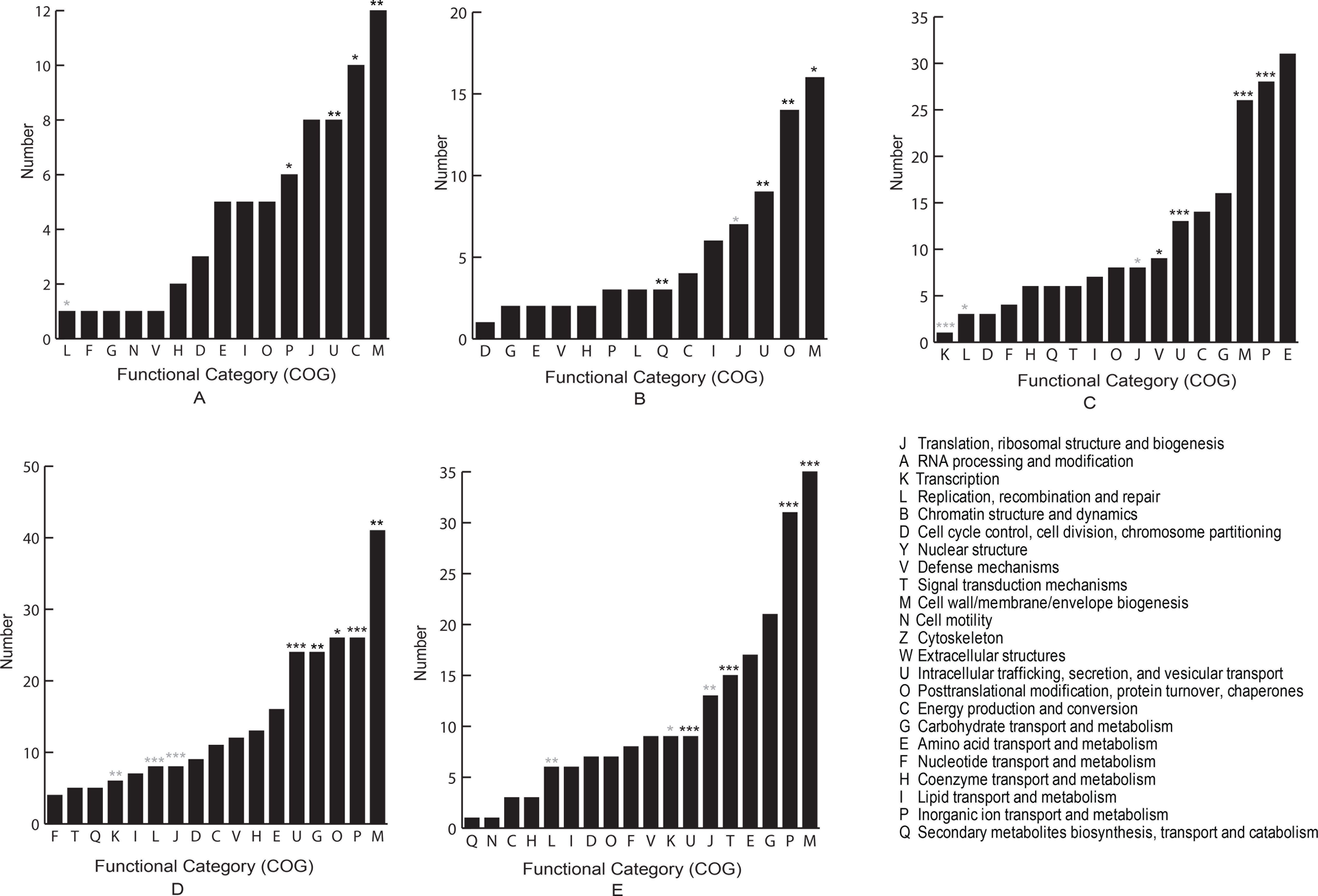
Analysis of Sec-dependent secreted proteins (secretome) of representative bacteria from clades with the strongest functional divergence. These clades are from SecA, SecB, SecE, SecG and SecY, respectively. Substrates are grouped functionally according to the classification of proteins into the Cluster of Orhtologous Groups (COG). These functional categories were ranked and plotted according to the numbers of their substrates. (A) *Chlamydia trachomatis*, (B) *Rickettsia prowazekii*, (C) *Acinetobacter baumannii*, (D) *Xylella fastidiosa*, (E) *Streptococcus agalactiae*. COG functional categories showing significant enrichment (black stars) or impoverishment (grey stars) of predicted substrate proteins are labelled by (*, P<0.05; **, P<0.01; ***, P<0.001).

**Figure 13.**
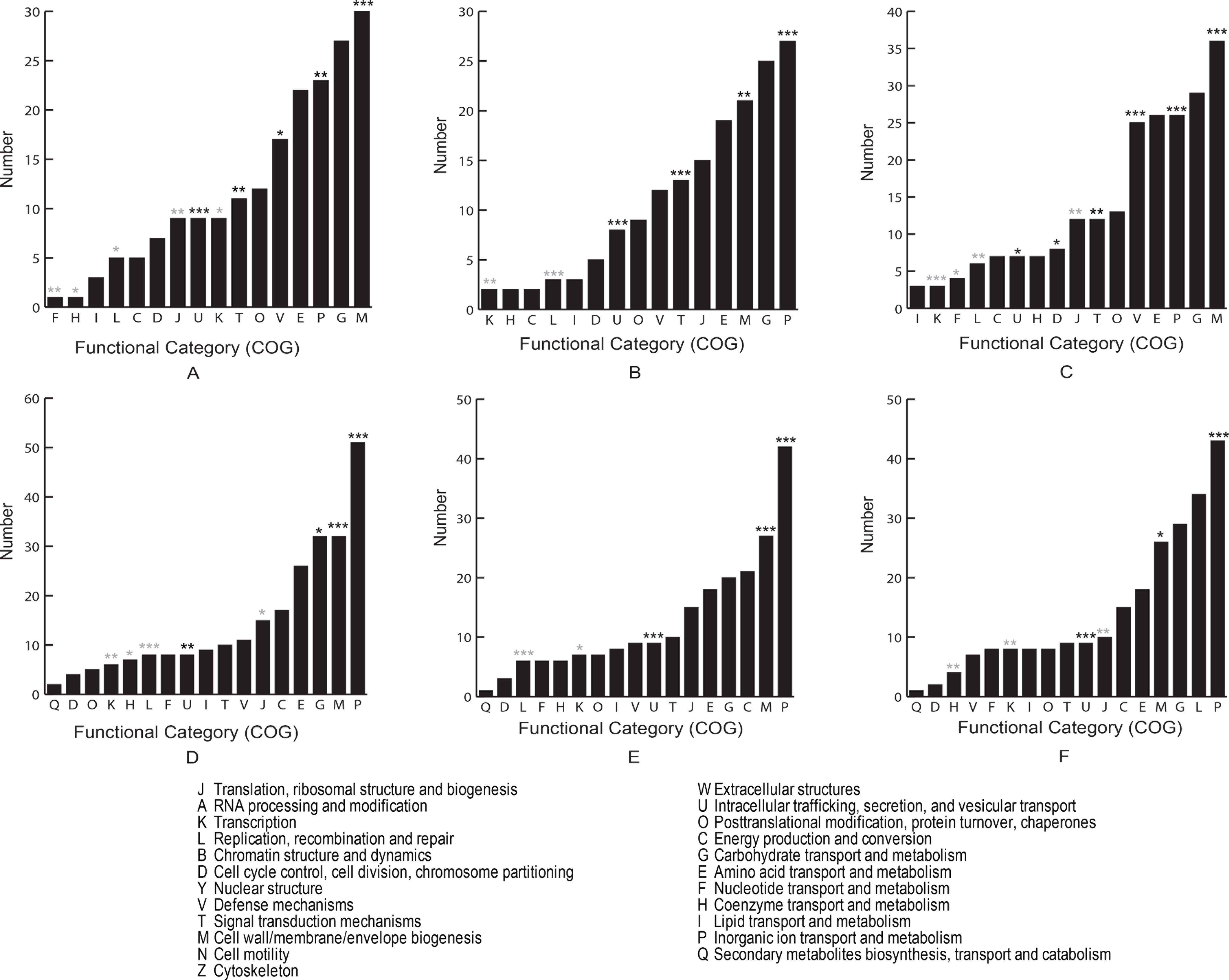
Analysis of Sec-dependent secreted proteins (secretome) from *Streptococcus* and *Staphylococcus*. Three species are chosen for *Streptococcus* and *Staphylococcus* from the clades with strong functional divergence, respectively. Substrates are grouped functionally according to the classification of proteins into the Cluster of Orhtologous Groups (COG). These functional categories were ranked and plotted according to the numbers of their substrates. (A) *Streptococcus gordonii*, (B) *Streptococcus pneumoniae*, (C) *Streptococcus sanguinis*, (D) *Staphylococcus aureus*, (E) *Staphylococcus epidermidis*, (F) *Staphylococcus haemolyticus*. COG functional categories showing significant enrichment (black stars) or impoverishment (grey stars) of predicted substrate proteins are labelled by (*:, P<0.05; **, P<0.01; ***, P<0.001).

## Discussion

In this study we demonstrated that the SecYEG-SecA system of pathogenic bacteria is the most functionally divergent among the bacterial lineages and that this FD has particularly followed the origination of SecA2/SecY2. Although this is expected, our study illuminates the question of how FD has occurred at the molecular level and how SecA/SecY functions have been affected and specialized in pathogenic bacteria. Importantly, FD has affected important domains in protein translocation. This evolutionary events may have mediated interaction of bacteria with their host, as previously suggested [18,19,21,22,25,26,29–32,49,68,87–96].

### Functional divergence in the ATP binding domain (NBD1) of motor protein ATPase SecA

We show that NBD1 and PPXD are the most enriched domains for sites under FD. NBD1 contains the ATPase activity, which is requied for the right conformational changes in SecA. These conformational changes are crucial in the transfer of the preprotein into the transmembrane SecY complex. PPXD seems to be involved in protein substrates intake and release and is believed to be the most dynamic domain, as supported by our structural analyses (Figure 3 and Table 1). Because of their fundamental role in the interaction with SecA substrates, their high enrichment for FD may have specialised SecA to interact with substrates specific to the ecological niche of the bacterium.

The first low micromolar inhibitors of bacterial SecA was recently synthesized following a preliminary *in silico* screening and have proven effective *in vitro* and *in vivo* against *E. coli* strains [64,70]. Authors of the two studies showed that the parent compound of the synthesized inhibitors form hydrogen bond specifically with a region located in the NBD1 domain of SecA (Figure 7E). They also showed that such a compound interacts with a residue (417Gly) located in the PPXD domain and a residue (534Arg) located in the NBD2 domain. The amino acid sites identified to have undergone FD in our study are in the vicinity of these regions (Table S5), hence being promising targets for novel inhibitory compounds. Moreover, NBD2 and HSD domains shared similar profiles of FD, probably due to the modulation of HSD domain function or structure by NBD2, possibly through the two-helix finger, proposed previously to participate in moving the substrate polypeptide chain into the SecY channel [69].

Importantly, we showed that NBD1 domain is the most functionally divergent domain in SecA from pathogenic bacteria. Similar molecular patterns of FD in the SecYEG-SecA protein complex were observed among major human and animal bacterial pathogens such as *Chlamydia, Streptococci, Staphylococci, Mycobacteria, Listeria, Legionnella* and *Mycoplasma*. *Chlamydial* species are obligate intracellular bacterial pathogens and have a unique lifestyle, requiring a special set of membrane proteins to interact with the host [97]. The secretion of these proteins may have been possible through FD changes in SecA of *Chlamydia*.

Amongst the most affected bacterial clades for FD was also *Coxiella burnettii*. The range of hosts that *Coxiella* and *Mycoplasma* can infect is very wide and includes arthropods, fish, birds, and a variety of mammals [98,99]. This might have required specific protein sets to invade a variety of ecological niches. We also found that the sequences of SecYEG-SecA of all the *Mycobacteria tuberculosis* strains so far sequenced are identical both at the protein and nucleotide levels and probably present similar pathogenic potential for Sec-dependent virulence. Interestingly, these bacteria have been proposed to be prone to acquire virulence factors by horizontal gene transfer [100], sparking speculation that gain and loss of Sec-dependent host colonization factors may contribute to host specificity, such as in *M. bovis* (host: cattle) and *M. tuberculosis* (host: human).

Some of the most affected pathogenic bacteria with FD are those that possess an accessory SecA2/SecY2 system and include highly pathogenic and antibiotic resistant bacteria such as *Staphylococcus aureus* (meticillin-resistant, vancomycin-susceptible, vancomycin-intermediate resistance strains) and *Streptococcus pneumoniae* etc. Although not essential for pathogens survival [18,19] this system is responsible for secreting a set of virulence factors [18]. It has been demonstrated that some gram-positive bacteria have an accessory SecA2/SecY2 system [18], which has been suggested to be involved in translocation of post-translationally modified (for example glycosylated) proteins in pathogenic *Streptococcus* and *Staphylococcus* species [20–29]. Given that glycosylated preproteins are large cell surface glycoprotein (about 2000-5000 amino acids,) we would expect FD to have taken place in SecA2 and SecY2 in comparison with the canonical proteins to cope with two important changes: greater conformational changes in SecA2 through an optimization of the regulation of ATP hydrolysis and interaction with SecY2 channel [30–32]; and SecY2 channel had to translocate preproteins with heavily glycosylated amino acid residues. Although we demonstrate that strong FD has occurred in key amino acid sites of SecA2 and SecY2 for ATP hydrolysis and translocation, further analyses are required to confirm the direct link between this FD and an optimized translocation of glycosylated proteins.

Clustering of bacteria according to FD has also brought forward some interesting results, among which we highlight the clustering of some *Mycobacteria* that lack SecY2 with a group consisting of seven plant species. Recent studies have shown that secretion of superoxide dismutase A (SodA) is SecA2 dependent, and SodA may help *M. tuberculosis* survive the oxidative attack of macrophages and thus plays a role in *M. tuberculosis* pathogenesis [32,87,101]. *M. tuberculosis* SodA belongs to the iron Sod (FeSOD) group, which has five homologues in *Arabidopsis thaliana*. It has been shown that two of the five homologues are FeSOD *(fsd2* and *fsd3*) and located in the chloroplast, where they play essential roles in early chloroplast development [102]. Such a remarkable convergent evolution between bacterial SecA2 and chloroplast SecA is plausibly the result of their adaptation to secreting similar sets of proteins.

### Radical functional divergence in SecA ATPase may compensate for the lack of SecB chaperone

Gram-positive pathogens seem to lack the molecular chaperone SecB (for example using *E. coli* SecB as query against KEGG SSDB database returns no significant hits in these bacteria). This is surprising since SecB mediates the post-translational translocation. Other proteins may therefore substitute SecB function, although these proteins remain elusive. The identification of a chaperone activity for SecA raises the possibility that this protein has a role in post-translational translocation [4]. We show that SecA protein from gram-positive bacteria has accumulated the most radical changes compared with other bacterial lineages supporting a different role of this protein in these bacteria. The fact that a mutation of *E. coli* SecA can partially compensate for the absence of SecB chaperone [3], points to that radical amino acid substitutions in SecA are likely to confer this protein the ability to bind preproteins and perform chaperone-like activities.

### Functional divergence and intermolecular coevolution of prokaryotic translocon channel and motor protein mediates ecological adaptation

The translocon core protein SecY (Sec61 in eukaryotes) is conserved within all three domains of life, supporting strong selective constraints on this protein within each of the domains. In addition, coevolutionary analysis on the accessory SecA2/SecY2 system indicates that there may be common mechanisms shared between the accessory and canonical SecA/SecY systems. For instance, the ATPase activity of SecA2 seems to be coupled with the channel opening and sealing of SecY2, which was evident in the canonical SecA/SecY system as well [103]. These coevolving sites may play an important role in maintaining the stability of the plug displacement upon SecA2 binding to SecY2 as suggested in the canonical SecA1/SecY1 system [12]. Given our results we propose four conditions to be met for the translocation of proteins to take place through this complex. First, amino acid changes in the ATPase SecA2 may be required to help the PPXD domain to effectively interact with the highly glycosylated substrates. Second, a wider channel opening state may also be required to cope with the glycosylated amino acid residues. Third, the NBD2 domain on SecA2 should play an important role in tuning a “translocation competent” state of SecY2 protein for translocating these large glycosylated cell surface proteins. Lastly, sites on the plug domain (p1), together with others located on other SecY2 protein domains (e.g. T7) should cope with the binding of the long N-terminal signal peptide of the pre-protein in a coordinated way and therefore their coevolution is essential.

### Functional divergence of the translocon channel core SecY contributes to the diversification of major proteobacteria groups

SecY is evolutionary conserved across all domains of life and plays a fundamental role in the biogenesis of membrane proteins and cell surface proteins of prokaryotes. Moreover, membrane proteins and cell surface proteins directly determine the lifestyle that a prokaryotic species can adopt, as they are at the interface of bacteria and environment [104–107]. Therefore, diversification of SecY function contributes to the diversification of major proteobacteria groups. Indeed, we show that the levels of FD in SecY differ greatly with the diversification of the major proteobacterial groups and link tightly with their ecological contexts.

## Conclusions

In this study, we revealed the main evolutionary forces that have driven the evolution of the SecA-SecYEG complex. We show that these proteins have diverged the most in pathogenic bacteria compared to non-pathogenic ones. In particular, SecA2 and SecY2 have diverged in function in pathogenic bacteria possibly to drive the secretion of a set of proteins specialised in pathogenesis. Finally, we unveil the molecular evolutionary mechanisms in these secretion proteins and their potential roles in niche specific adaptations. Because of their importance in the ecological adaptation of bacteria, and in particular of pathogenic bacteria, we propose these proteins and important amino acid sites as possible drug targets for future therapeutic drugs.

## Acknowledgments

This work was supported by Science Foundation Ireland, under the program of the President of Ireland Young Researcher Award to Mario A. Fares, and the IRCSET (Irish Research Council for Science, Engineering and Technology) Government of Ireland Postgraduate Scholarship in Science, Engineering and Technology to Xiaowei Jiang. This work was also supported by grant BFU2009-12022 funded by the Spanish Ministerio de Ciencia e Innovatión to MAF. The funders had no role in study design, data collection and analysis, decision to publish, or preparation of the manuscript. We would like to thank some members of the Fares’ lab for a critical reading of the manuscript.

## Author contributions

Conceived and designed the experiments: MFA and XJ. Analyzed the data: XJ. Drafted the manuscript XJ. Wrote the final version of the manuscript MAF.

## Supplementary Tables

**Table S1.** Species in all clades under functional divergence annotated with IDC-10 for SecA. All the clades are ranked according to their number of sites detected under functional divergence.

**Table S2.** Species in all clades under functional divergence annotated with pathogenic status for SecB. All the clades are ranked according to their number of sites detected under functional divergence

**Table S3.** Species in all clades under functional divergence annotated with IDC-10 for SecE. All the clades are ranked according to their number of sites detected under functional divergence.

**Table S4.** Species in all clades under functional divergence annotated with IDC-10 for SecG. All the clades are ranked according to their number of sites detected under functional divergence.

**Table S5.** Species in all clades under functional divergence annotated with IDC-10 for SecY. All the clades are ranked according to their number of sites detected under functional divergence.

**Table S6.** Secretome analysis of 207 bacterial genomes. This table includes all COG functional categories that have been statistically tested to be over-enriched with three levels of P-values: p<0.05; p<0.01; p<0.001. Clades are also grouped to their closest phylogenetic group.

**Table S7.** The mean amino acid atomic distance between amino acid sites that have been identified to be inhibitor binding in citation [64] and amino acid sites identified to be under FD in this study. Sites are located in SecA NBD1 domain, and the protein structure used for calculating the atomic distance is 3DIN chain A.

## Supplementary Figures

**Figure S1.** Two-dimensional clustering of protein domains and all species under functional divergence for SecA.

**Figure S2.** Two-dimensional clustering of protein domains and all species under functional divergence for SecY.

**Figure S3.** Networks of coevolving sites between SecA, SecB and SecY. Residues in the networks are sorted clockwise in ascending order depending on the number of coevolutionary interactions each amino acid residue establishes. Properties of amino acid sites are explained as follows. Amino Acid: amino acid sites are mapped to *E. coli* SecA. Degree: number of coevolving partners. Domain: protein domain in which a coevolving site locates. Protein: the protein where the coevolving sites locate.

